# Insulin-degrading enzyme activity is modulated by interaction with pyrroline-5-carboxylate reductase 1

**DOI:** 10.64898/2026.07.16.739082

**Authors:** Eun Suk Song, Camila Camacho-Navas, Anwesha Goswami, Anmol Nayak, Karol A. Arizaca Maquera, Jing Chen, Stefan Stamm, Emilia Galperin, Louis B. Hersh, David W. Rodgers

## Abstract

Insulin-degrading enzyme (IDE, insulysin, insulinase) is a peptidase that hydrolyzes a number of bioactive peptides including insulin and the amyloid beta peptide, making it a promising therapeutic target for diabetes and Alzheimer’s disease. Aspects of its physiological role are still controversial, however. In an attempt to further define IDE’s role in cells, we used co-immunoprecipitation experiments to identify potential IDE interacting proteins. The enzyme pyrroline-5-carboxylate reductase 1 (PYCR1) was found associated with IDE in three different cell lines, and the two proteins colocalize in HeLa cells. Purified PYCR1 activates IDE toward small peptide substrates, suggesting a modulatory role for the interaction *in vivo*. Modeling suggests that the unstructured N-terminal region of PYCR1 inserts into allosteric sites of IDE, contributing to the observed activation. Deleting this sequence alters, but does eliminate, the interaction between PYCR1 and IDE. Since pyrroline-5-carboxylate reductase 1 is a mitochondrial protein, we posit that their interaction could regulate a previously described mitochondrial pool of IDE, which may serve to degrade mitochondrial targeting sequences or amyloid beta peptide that localizes to that organelle.

## Introduction

Despite having been described more than 70 years ago [1, 2] the physiological roles of insulin-degrading enzyme (IDE, insulinase, insulysin) have not been well defined [3]. There is considerable evidence supporting IDE’s role in insulin and amyloid beta peptide metabolism [4–6]. However, these functions have been questioned [7–9], and roles for IDE in other physiological processes have been proposed, including serving as a receptor for Varicella Zoster Virus [10], stimulating insulin secretion [11, 12], playing a role in ubiquitinated protein clearance [13], and acting as a heat shock protein [13].

IDE likely functions as a homodimer in which each subunit is composed of four domains. The overall structure [14] resembles a clamshell, which is believed to open and close during the catalytic cycle. The open form would bind its peptide substrate in a large inner chamber of the enzyme while the closed form would catalyze the cleavage of the peptide substrate. A conformational change back to the open form would then allow the release of products. The subunits in the IDE dimer can function independently, but there can also be communication between subunits in which substrate binding to one subunit increases the turnover of the adjacent subunit [15]. Kinetic studies have shown that IDE exhibits allosteric activation in that the hydrolysis rate of synthetic substrates and small peptides such as angiotensin and bradykinin can be increased 5 to 100-fold by polyphosphates, while larger peptides such as insulin and glucagon are largely unaffected [16, 17]. This activating effect results from the interaction of the activator with an anion binding region formed by the inner surface of the IDE C-terminal domain [16].

In addition to its many proposed functions, multiple subcellular locations for IDE have been described. The majority of the enzyme is in the cytosol [18, 19], but some of the intracellular pool localizes to endosomal compartments where it can degrade target peptides [3, 4, 20–22]. Also, a minor IDE isoform that includes an additional 41 N-terminal amino acids has been reported to localize to mitochondria [23–26], and IDE trafficking to peroxisomes [27] as well as the plasma membrane [28] has been reported. A number of studies have proposed that IDE is secreted from cells by a mechanism insensitive to classical Golgi-based secretion inhibitors [29–31], but these claims have been disputed [3, 8]. Our finding that IDE in the media of several cultured cell lines is proportional to the amount of the known intracellular protein lactate dehydrogenase supports the conclusion that extracellular IDE is the result of cell breakage and not secretion [32].

It has been suggested that a search for potential interaction partners of IDE might provide important insight into its true functions, regulation, and localization [3]. However, few studies looking for IDE binding partners have been reported. IDE has been proposed to interact with steroid receptors [33] as well as the proteasome [34, 35], and it was found to bind disassociated nestin/vimentin complex leading to inhibition of insulin-degrading activity and an enhancement of small peptide cleavage [36]. Here we report the results of a study in which a specific anti-IDE nanobody was developed and used in co-immunoprecipitation (Co-IP) experiments to identify the mitochondrial enzyme pyrroline-5-carboxylate reductase 1 (PYCR1) as an IDE interacting and potentially regulating protein.

## Materials and methods

### Cultured cells and IDE production

HEK293 and COS-1 cells were obtained from ATCC (Manassas, VA). Cell identities were verified by the vendor. HeLa cells and IDE knockout HeLa cells were obtained from Abcam (ab261755) and verified by the vendor. The IDE knockout cells were shown by the vendor not to express full-length IDE by Western blotting, and we confirmed this finding. Furthermore we found IDE knockout HeLa cells lacked IDE activity. Cells were cultured as described previously [32]. Recombinant IDE and the N-terminal half of IDE were produced as described [37].

### Production of an anti-IDE nanobody

Nanobodies to IDE were produced by immunizing an alpaca with purified rat IDE according to Chow et al. [38]. One of these nanobodies, nanobody D1, which had a high affinity for IDE (see below), was purified to homogeneity by nickel affinity chromatography followed by molecular sieve chromatography on a Cytiva Superdex S-75 column.

The binding affinities of the nanobodies for IDE were determined by bio-layer interferometry using a Blitz instrument (Sartorius) according to established protocols [39]. A nanobody bearing a polyhistidine tag was immobilized on an Anti-Penta-His biosensor (Sartorius 18-5101) and washed with 50 mM Tris (pH 7.4). IDE on and off rates were determined in the same buffer. Rate constants (k_on_ and k_off_) and affinity (K_d_) were determined in GraphPad Prism 7.0 using the association then dissociation model (Fig S1). Anti-IDE nanobody D1 had a K_d_ for intact IDE of 473 nM (95% CI: 451-484 nM). This same nanobody bound the N-terminal half of IDE with a K_d_ of 421 nM (95% CI: 401-441 nM). As expected, a negative control nanobody raised against viral P protein (C9) had a more than 1000-fold lower affinity for intact IDE of ∼70 mM.

For Co-IP assays, the Fc portion of rabbit IgG was inserted at the 5’ end of the coding sequence of the D1 nanobody (Fc-anti-IDE-Nb-D1) in the expression vector pFUSE-rlgG-Fc-2 (pfuse-rfc-2: InvivoGen) to allow interaction with protein A beads. A similar construct was prepared using the C9 control nanobody (Fc-anti-PP-Nb-C9). Expi293 cultured cells (ThermoFisher) were used to produce the constructs, which were purified with a Hi Trap Protein G HP column (Cytiva).

### Co-immunoprecipitation assays for interacting proteins

Proteins interacting with IDE were immunoprecipitated from cellular extracts with the Fc-anti-IDE-Nb-D1 nanobody fusion bound to protein A beads. The Fc-anti-PP-Nb-C9 nanobody fusion was used as a negative control. The protein A beads (Abcam 193256, ∼40 μl) were washed with IP buffer (50 mM Tris pH 7.4, 150 mM NaCl, 1% Triton X-100, 1 mM Na_3_VO_4_, 10 mM NaF) containing protease inhibitors without EDTA (Sigma-Aldrich), added to 50 μg of nanobody, and then incubated for ∼2 h on a rotator at 4°C. After incubation, samples were centrifuged for 5 min at 500 g and 4°C to collect the beads.

Cellular extracts were prepared in IP buffer containing protease inhibitors without EDTA (Sigma-Aldrich) by gentle rocking for 45 min at 4°C, and clarified lysates were obtained by centrifugation for 20 min at 14,000 × g. The clarified lysates were then adjusted to ∼2 mg/ml protein. Eighty microliters of nanobody bound protein A agarose beads was added to 5 ml of extract (total of 10 mg of lysate protein) and incubated for 2 hrs. on a rotator at 4°C. The protein–antibody bead complex was then washed 6 times with IP buffer and the beads collected by centrifugation. Bound proteins were released from beads by adding 80 μl of 2×SDS sample buffer. Samples were then loaded and run on Sure PAGE, Bis-Tris 4-12% gels (GenScript). Gels were stained with Coomassie blue, and bands were excised from the HEK 293 cell gel and identified by mass spectrometry of tryptic peptides. For the COS-1 and HeLa cells, pyrroline-5-carboxylate reductase 1 (PYCR1) and glucose-6-phosphate 1-dehydrogenase (G6PD) were identified by Western blots using specific antibodies (G6PD – Bethyl Laboratories A300-404A, PYCR1 – Proteintech 13108-1-AP).

### Identification of interacting proteins by mass spectrometry (LC-ESI-MS/MS) analysis

All mass spectra reported in this study were acquired by the University of Kentucky Proteomics Core Facility. Excised gel bands were subjected to dithiothreitol reduction, iodoacetamide alkylation, and in-gel trypsin digestion as previously described [40]. The resulting tryptic peptides were extracted, concentrated, and subjected to shot-gun proteomics analysis. LC-MS/MS analysis was performed using an LTQ-Orbitrap mass spectrometer (ThermoFisher Scientific, Waltham, MA) coupled with an Eksigent Nanoflex cHiPLC™ system (Eksigent, Dublin, CA) through a nano-electrospray ionization source. The peptide samples were separated with a reversed phase cHiPLC column (75 μm x 150 mm) at a flow rate of 300 nL/min. The mobile phase A was water with 0.1% (*v*/*v*) formic acid while the mobile phase B was acetonitrile with 0.1% (*v*/*v*) formic acid. A 50 min gradient condition was applied: initial 3% mobile phase B was increased linearly to 40% in 24 min and further to 85% and 95% for 5 min each before it was decreased to 3% and re-equilibrated. The mass analysis method consisted of one segment with 11 scan events. The first scan event was an Orbitrap MS scan (300-1800 m/z) with 60,000 resolutions for parent ions followed by data dependent MS/MS for fragmentation of the 10 most intense multiple charged ions with the collision induced dissociation method.

The LC-MS/MS data were submitted to a local Mascot server for MS/MS protein identification via Proteome Discoverer (version 1.3) against a custom database containing reviewed Homo sapiens (Human) sequences from UniProt (total of 20,386 proteins). Typical parameters used in the Mascot MS/MS ion search were trypsin digestion with a maximum of two miscleavages, cysteine carbamidomethylation and methionine oxidation, and 10 ppm precursor ion and 0.8 Da fragment ion mass tolerances. A decoy database was built and searched. Filter settings that determine false discovery rates (FDR) were used to distribute the confidence indicators for the peptide matches. Peptide matches that pass the filter associated with the FDR rate of 1% and 5% were assigned as high and medium confident peptides, respectively.

### IDE activity

Activity toward the synthetic fluorogenic substrate Abz-Gly-Gly-Phe-Leu-Arg-Lys-His-Gly-Gln-EDDnp (Abz substrate) was determined as previously described [41] by following the increase in fluorescence upon peptide cleavage. Insulin and bradykinin cleavage were determined by following the IDE dependent decrease in their peak area as determined by HPLC [41].

### Immunofluorescence microscopy and analysis

HeLa wild type, and IDE-KO HeLa cells (Abcam) were grown on collagen-coated coverslips in DMEM media and transfected with a pcDNA-3.1 plasmid expressing full-length IDE (residues 1-1019, IDE-Met^1^) with a C-terminal HA tag. Prior to fixation, cells were incubated with MitoView™ Fix 640 dye (Biotium) for 2 h at 37°C in 5% CO_2_ followed by permeabilization with 0.5% saponin for 5 min. Cells were then fixed with 4% paraformaldehyde (Electron Microscopy Sciences) for 15 min at room temperature. Following fixation, cells were washed in PBS and blocked with 10% normal serum, 1% BSA for 1 h. For immunostaining, cells were incubated with primary antibodies (mouse anti-HA, Cell Signaling Technology 2367S; rabbit anti-PYCR1, Proteintech 13108-1-AP) for 90 min in PBS containing 1% BSA, rinsed, and incubated for 60 min with respective secondary antibodies (Alexa Fluor 488-or Alexa Fluor 647-conjugated; Invitrogen AB150073, AB150105, or AB150115) in PBS with 1% BSA at room temperature. Cells were rinsed and mounted for microscopy in Prolong mounting medium (Fisher Scientific).

All images were acquired using a Mariannas Imaging system consisting of a Zeiss inverted microscope equipped with a cooled CCD CoolSnap HQ (Roper), dual filter wheels, and a 175W xenon light source, all controlled by SlideBook software (Intelligent Imaging Innovations). The detection of DAPI stain, Alexa Fluor 488, Alexa Fluor 549 and MitoView™ Fix 640 fluorescence were performed using blue, green, TRITC and Cy5.5 channels respectively. Images were acquired in 2X2 binning mode, and image analysis was performed using deconvolution of Z-stack image sets with SlideBook image processing modules. Analyses of colocalization was conducted using the JACoP plugin [42] and Colocalization Threshold utility in Fiji [43], with intensity thresholds determined according to Costes [44]. Statistics for colocalization pixel overlaps are averages from three experiments over a total of 12 cells for IDE and mitochondria, 21 cells for PYCR1 and mitochondria, and 23 cells for IDE and PYCR1. Pixel size for all images represented 0.196 μm on the samples.

### Production of PYCR1

A PYCR1 cDNA was obtained from Addgene (a gift to Addgene by Nicola Burgess-Brown), which covered residues 1-300 of the 319 amino acid protein. A His tag coding sequence was added to the 5’ end of the cDNA, which was then expressed in *E. coli* BL21(DE). Recombinant PYCR1 was purified by nickel affinity chromatography followed by gel filtration. Two peaks of PYRC1 protein were observed from the gel filtration column, one corresponding to ∼400 kDa and the other corresponding to ∼200 kDa. Both peaks gave a ∼33 kDa monomer according to SDS-PAGE in agreement with the reported monomeric molecular weight [45].

Full length PYCR1 was obtained by synthesizing a C-terminus coding 149 base pair fragment which contained Nco1 and BamH1 sites and cloning this fragment into ^-^PYRC1^300^ using an internal Nco1 and a vector BamH1 site. A truncated PYCR1 (residues 1-275) was produced by synthesizing a fragment which contained KpnI and BamHI sites and cloning into the pNIC28-Bsa4 vector.

### PYCR1 activity and its direct interaction with IDE

The activity of PYCR1 was determined fluorometrically using the assay described by Meng et al. [46] in which the enzyme catalyzes the oxidation of thioproline with the concomitant reduction of NADP. The reaction was linearly dependent on PYRC1 as well as dependent on the presence of thioproline.

Biolayer interferometry was used to determine if IDE and PYCR1 interact directly. The Blitz instrument was employed largely as described above. PYCR1 protein with its N-terminal polyhistidine sequence was immobilized on HIS2 biosensors (Sartorius 18-5114) and washed with 20 mM Tris (pH 7.4), 150 mM NaCl. IDE on and off rates were then determined in the same buffer at IDE concentrations of 2.1, 4.2, and 8.3 µM. Biosensors were washed in 20 mM Tris (pH 7.4), 150 mM NaCl, 20 mM imidazole, 0.6 M sucrose, and 1% bovine serum albumin [47] for at least 10 minutes and then equilibrated into the original buffer between different IDE concentrations. Three replicates for each IDE concentration from separate experiments were including in the analysis, which was done using the association then dissociation model in GraphPad Prism. Global fitting of on and off rates across different concentrations was carried out by unweighted least squares regression.

Dynamic light scattering [48, 49] was used to determine the hydrodynamic radius and size distribution of the IDE, PYCR1, and the potential complex using a DynaPro NanoStar instrument (Waters Wyatt Technology). Samples were centrifuged at 16,000 ×g for 8 minutes to remove particles and aggregates and placed in a 2 µl sample volume quartz cuvette. Scattering data were collected at 20°C, and each measurement consisted of 10 acquisitions with 5-second integration times. Protein concentrations were: IDE 26.0 µM; PYCR1 47 µM; complex IDE 39 µM, PYCR1 31 µM. Data analysis was performed using DYNAMICS software, version 7.1.8.93, with autocorrelation functions fit and analyzed by the Dynals regularization algorithm, which provided size distributions, polydispersity estimates, and the hydrodynamic radius based on the Stokes–Einstein equation. The globular protein model was used to estimate molecular weights from the hydrodynamic radius.

### RNA isolation from HeLa cells

RNA was isolated [50] from HeLa cells using the RNeasy Micro Kit (Qiagen) following instructions by the manufacturer. RT-PCR was used for cDNA synthesis according to standard protocols [51] using two primer pairs: Primer pair 1 (forward primer AGAGGCCACGGGGCTATAC and reverse primer CGGACATTTTCTGGTCTGAG), which will produce a 151 nt product and primer pair 2 (forward primer TGTAACCCTTCCTCGCCTTA and reverse primer AACCAGCTGACTTGGAAGGA), which will produce a 178 nt product. For these primers the T_m_ was 65°C and the extension time was 45 sec. Samples were run in a 2% agarose gel for 55 min.

## Results

### Co-immunoprecipitation identifies potential IDE binding partners

To identify cellular proteins that interact with IDE, we conducted co-immunoprecipitation (Co-IP) experiments by incubating IDE with cellular extracts and using a rabbit anti-IDE-Fc nanobody fusion (Fc-anti-IDE-Nb-D1), or a control nanobody against a viral P-protein (Fc-anti-PP-Nb-C9). Protein A was used to bring down the Fc-nanobody-IDE complex and any bound proteins.

Co-IP of an extract from HEK 293 cells using Fc-anti-IDE-Nb-D1 identified several proteins that differed compared to proteins pulled down with the Fc-anti-PP-Nb-C9 control nanobody, or in the absence of nanobody (**Fig 1)**. Immunoprecipitated proteins were identified by mass spectrometry (S1 **Table)**.

**Fig 1.**
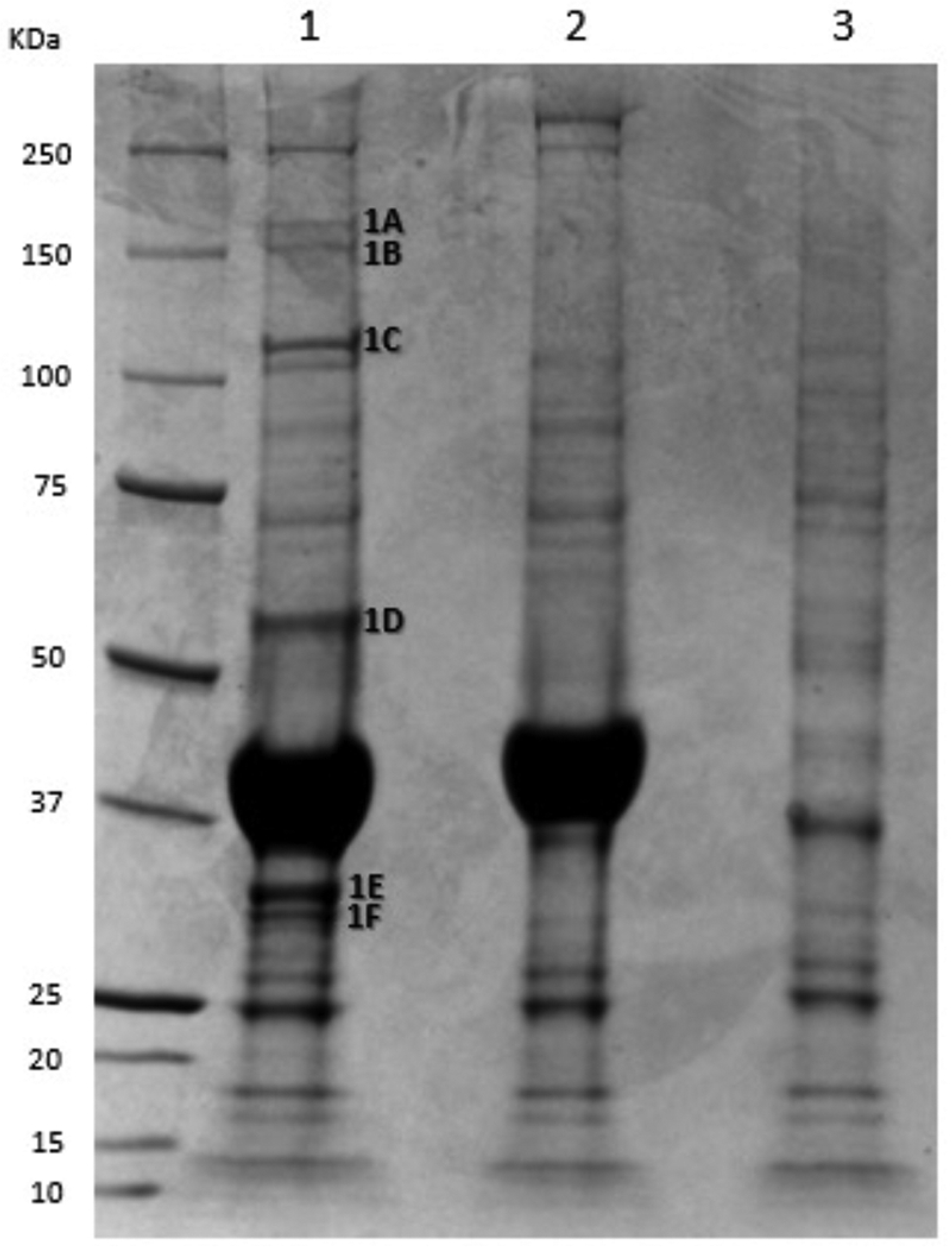
Co-immunoprecipitation from HEK 293 cells. An SDS-Page gel from a Co-IP of HEK 293 cellular proteins incubated with 1 - Fc-anti-IDE-Nb-D1, 2 - control nanobody Fc-anti-PP-Nb-C9, and 3- no nanobody. Each sample contained 100 µg of nanobody, 80 µl of washed Protein A, and 10 mg of HEK 293 cell lysate. Identified proteins unique to the IDE pulldown are: 1A = Kinesin-like protein KIF15 (Uniprot Q9NS87), 1B = Elongator complex protein (Uniprot O95163), Peroxisome biogenesis factor 1 (Uniprot O43933), 1C = Insulin-degrading enzyme, 1D = Glucose-6-phosphate 1-dehydrogenase (Uniprot P11413) and Vimentin (Uniprot P08670), 1E = Pyrroline-5-carboxylate reductase 1 (Uniprot P32322), Pyrroline-5-carboxylate reductase 2 (Uniprot Q96C36) and F = Purine nucleoside phosphorylase (Uniprot P00491).

To further establish the validity of the proteins that were identified in HEK 293 cells, we performed additional co-IP experiments using COS-1 and HeLa cells, both of which express IDE (**Fig 2**). Western blots confirmed that the two most abundant associated proteins from the HEK 293 line, pyrroline-5-carboxylate reductase 1 (PYCR1) and glucose-6-phosphate 1-dehydrogenase (G6PD), were also co-immunoprecipitated from the COS-1 and HeLa cells. In addition, comparison of the Fc-anti-IDE-Nb-D1 Co-IP gel lanes from the three cell lines indicated that all gel bands identified by mass spectrometry for the HEK 293 cells were also present in the two other lanes (**Fig S2**). Consistently finding the proteins in immunoprecipitants from three different cell lines increased confidence that they might be physiologically significant partners of IDE.

**Fig 2.**
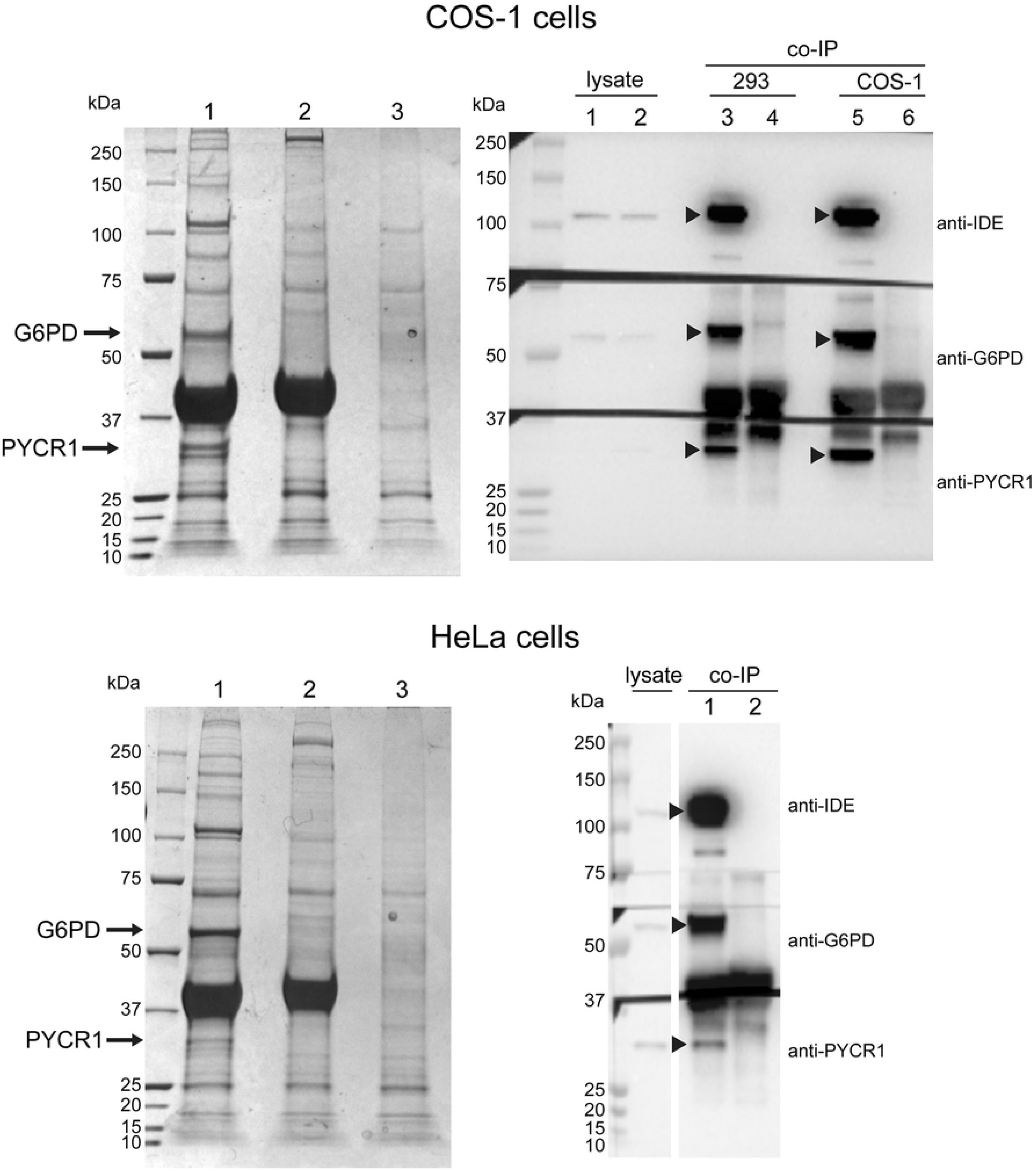
Identification of PYCR1 and G6PD in HeLa and COS-1 extracts. Cell extracts derived from COS-1 cells (top) and HeLa cells (bottom) were subjected to co-immunoprecipitation. Left panels show Coomassie-stained SDS-PAGE gels of the immunoprecipitants using (1) anti-IDE nanobody Fc-anti-IDE-Nb-D1, (2) control nanobody Fc-anti-PP-Nb-C9, and (3) the absence of a nanobody as described in Methods. Right panels show Western blots with the primary antibodies indicated. The blot for the COS-1 cells included an HEK 293 cell immunoprecipitant for comparison. Lanes are (1) 293 cell lysate, (2) COS-1 cell lysate, (3) HEK 293 IP with anti-IDE nanobody Fc-anti-IDE-Nb-D1, (4) HEK 293 IP with control nanobody Fc-anti-PP-Nb-C9, (5) COS-1 IP with anti-IDE nanobody Fc-anti-IDE-Nb-D1, (6) COS-1 IP with control nanobody Fc-anti-PP-Nb-C9. For the HeLa cells, the lysate is shown and along with IPs using (1) anti-IDE nanobody Fc-anti-IDE-Nb-D1, and (2) control nanobody Fc-anti-PP-Nb-C9. Arrowheads on the blots indicate IDE (top), G6PD (middle), and PYCR-1 (bottom). Molecular weight markers are indicated.

To further confirm the validity of the proteins co-immunoprecipitating with the IDE nanobody, we compared proteins immunoprecipitated from HeLa cells to proteins immunoprecipitated from HeLa IDE knock-out (KO) cells, which are devoid of both IDE activity as well as full-length IDE protein. In the HeLa KO cells, the IDE gene is disrupted by the introduction of a 4-base deletion in exon 14 of IDE (Abcam ab261755). Unexpectedly, the candidate IDE interacting proteins, PYCR1 and G6PD, were co-immunoprecipitated from the HeLa KO cells by nanobody Fc-anti-IDE-Nb-D1, but not by the control nanobody (**Fig S3)**. We considered two explanations for this finding: either the Fc-anti-IDE-Nb-D1 nanobody interacted with both G6PD and PYCR1, or the N-terminal part of IDE was still expressed and interacted with these proteins.

We focused our studies on PYCR1 as it was one of the most abundant pulldown proteins, and we were able to easily produce the recombinant protein using expression constructs that were readily available. An ELISA assay showed that the D1 nanobody, while clearly binding to IDE, did not bind to recombinant PYCR1. However, we noted that the four-base deletion in exon 14 of IDE could potentially produce an mRNA encoding the N-terminal region of IDE with an extension into the C-terminal half of the molecule (**Fig 3**). The predicted protein would be similar to the N-terminal half of IDE that we previously expressed [37]. Furthermore, using biolayer interferometry, the D1 nanobody was shown to bind to the N-terminal half of IDE (see **Fig S1)**, indicating that this IDE fragment could be immunoprecipitated in the pulldown assay. The IDE gene disrupting insertion is right after the codon for S557, which starts the second strand in the IDE domain 3 central sheet, and 54 potentially translated codons follow the S557 codon before a stop codon is reached (see **Fig 3**).

**Fig 3.**
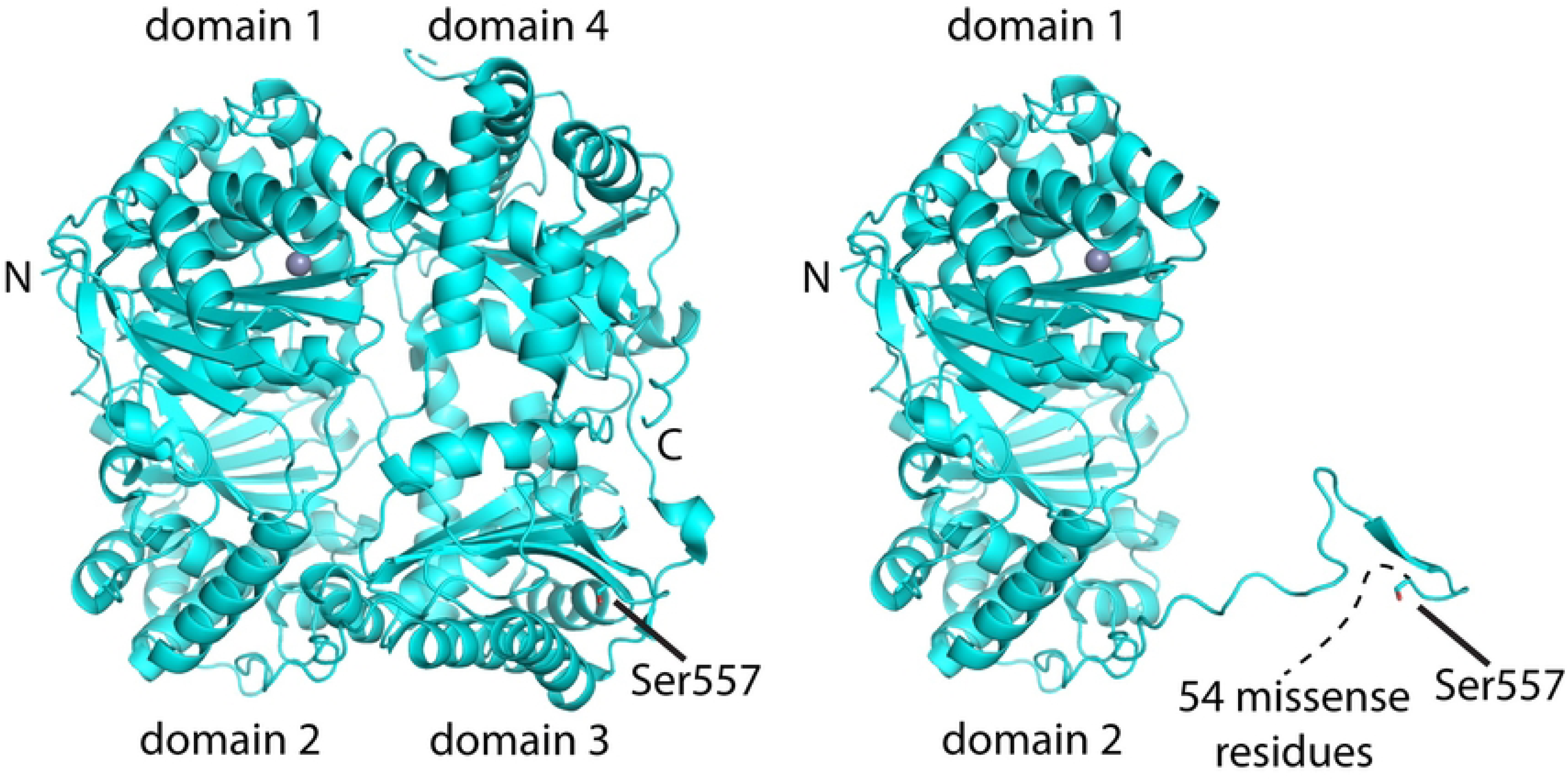
IDE fragment made in knockout HeLa cells. Left: Structure of wild type IDE. Right: Possible structure of the IDE fragment produced in HeLa IDE-KO cells.

To provide evidence that in fact this IDE fragment is produced in the HeLa IDE KO cells, we isolated RNA from both wildtype and IDE-KO cells and generated cDNAs using two specific primer pairs. As shown in **Fig 4**, both wild type HeLa cells as well as the HeLa IDE KO cells, yielded the same expected PCR products, showing that the HeLa IDE KO cells produced an mRNA consistent with translation of the N-terminal IDE fragment. In addition, a lysate from the HeLa IDE KO cells gave a band by Western blotting that was consistent with the expected size (60-65 kDa) of the IDE N-terminal fragment (Fig S4). (Smaller Western positive fragments, likely proteolytic products, are also seen.) It therefore seems likely that the proteins pulled down with IDE interact with the N-terminal half of the molecule. In that light, it is interesting to note that dimerization of IDE occurs via the C-terminal half of the molecule [14], which would make it less accessible to interacting proteins.

**Fig 4.**
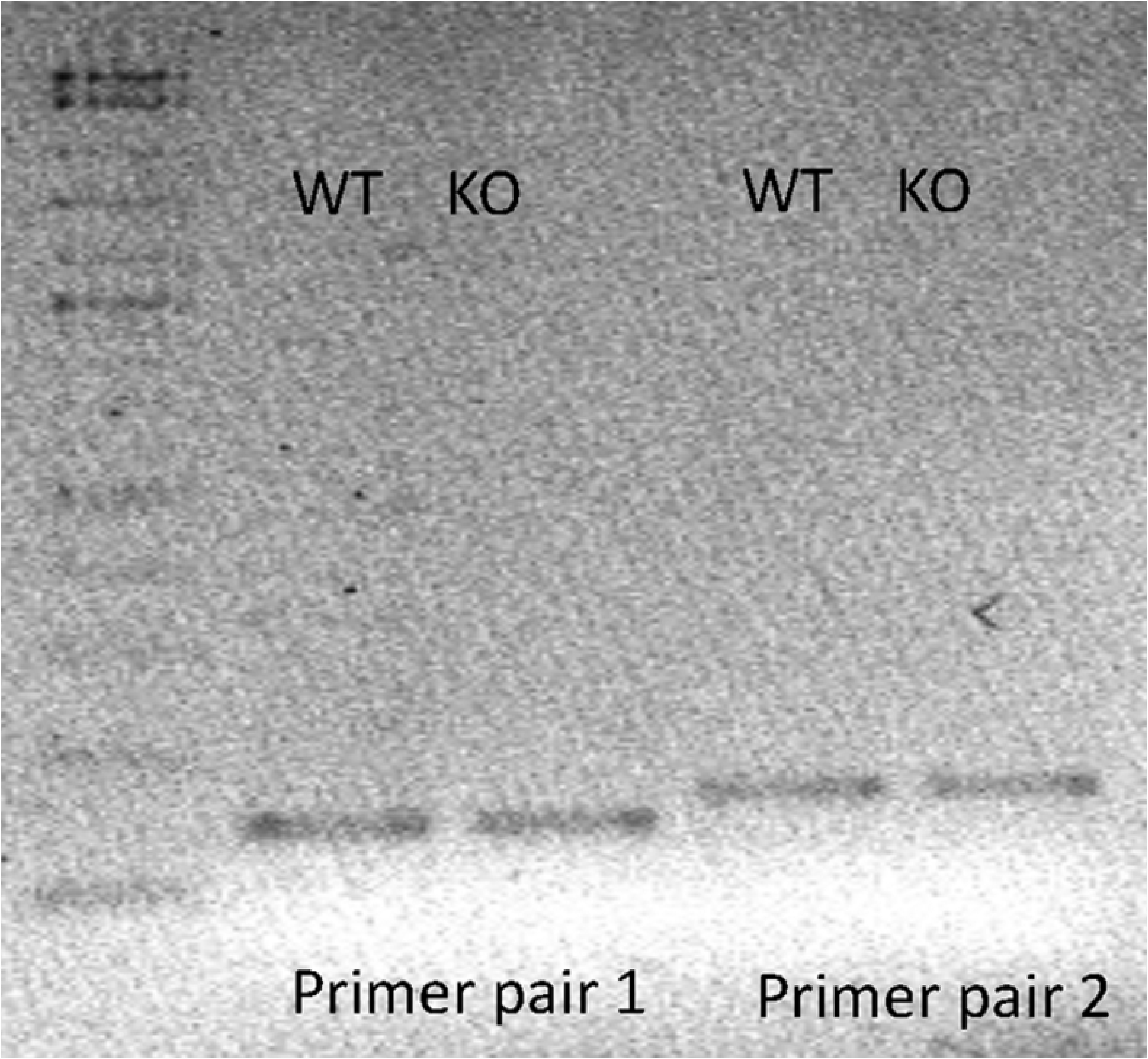
Wild type and HeLa-KO cells produce RNA fragments of the same apparent size. RNA was isolated from both wild type and IDE-KO HeLa cells as described in methods. Results of PCR reactions using the two primer pairs are shown, with the same size IDE RNA fragments generated from both wild type and IDE-KO Hela cells.

### IDE colocalizes with PYCR1 in mitochondria

Nuclear encoded PYCR1 localizes to mitochondria [52, 53], while the primary isoform of IDE produced in cells (residues 42-1019) is largely cytosolic [18, 19]. It has been shown, however, that the longest isoform of IDE (residues 1-1019, IDE-Met^1^) traffics to mitochondria [23]. We therefore expressed IDE-Met^1^ in IDE-knockout HeLa cells and determined its localization, along with PYCR1, by immunofluorescence microscopy. Both IDE-Met^1^ and PYCR1 colocalized with mitochondria (**Fig 5 and Table 1**), giving Manders’ colocalization coefficients [54] of 0.91 and 0.76, respectively, for pixel overlap relative to the mitochondrial marker above threshold. IDE-Met^1^ and PYCR1 also showed extensive colocalization, with Manders’ coefficients of 0.88 and 0.78, respectively, for pixel overlap of their intensities and a Pearson’s correlation coefficient [55] of 0.69. Co-IP of PYCR1 may have resulted from partial permeabilization of mitochondria and interaction with cytosolic IDE, but colocalization of IDE-Met^1^ and PYCR1 suggests that their physiologically relevant interaction occurs in mitochondria.

**Fig 5.**
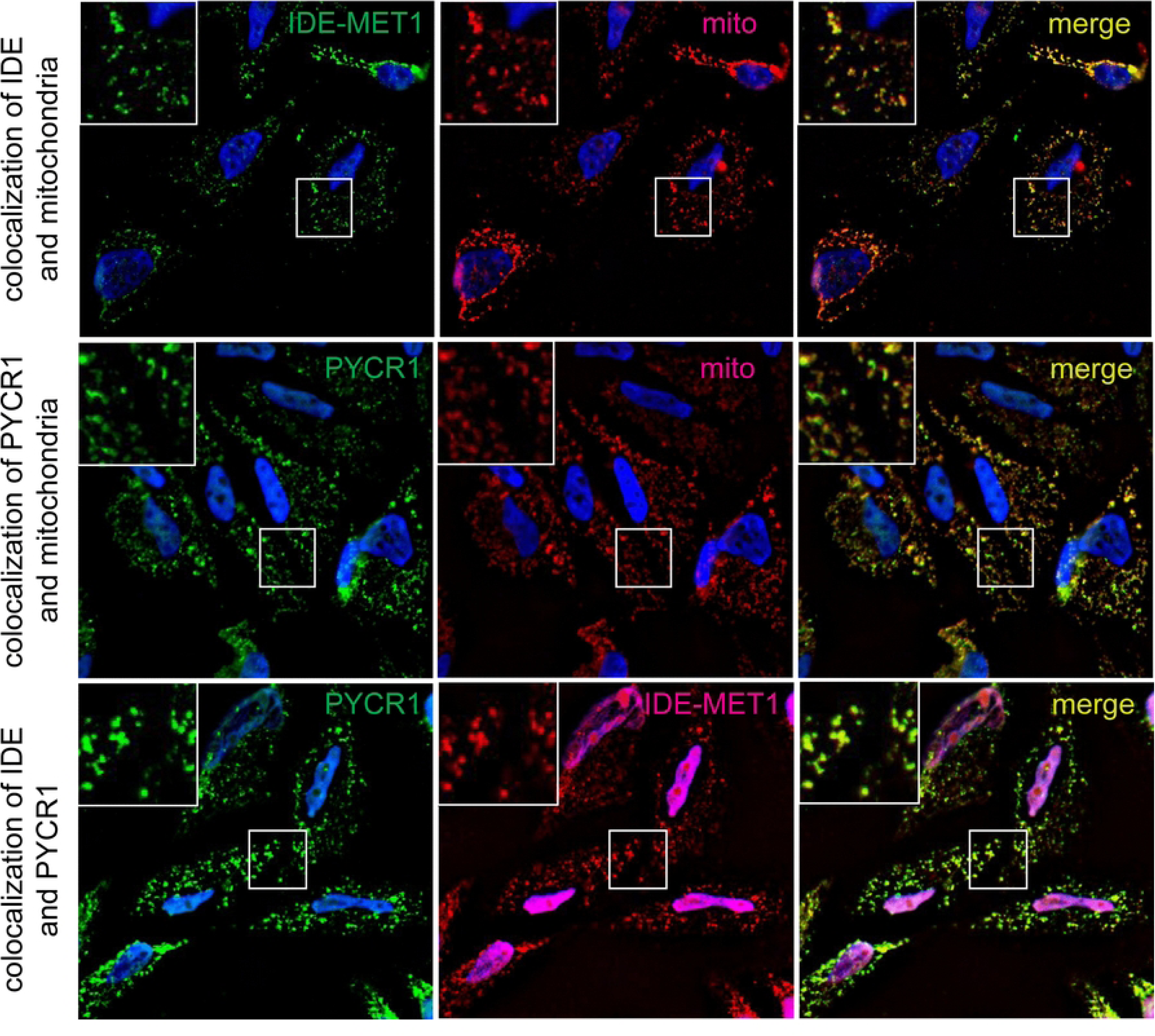
Colocalization of IDE and PYCR1 in mitochondria of IDE knockout HeLa cells. Immunohistochemistry fluorescent microscopy showing colocalization of IDE and PYCR1 in mitochondria. Top row: Cells expressing the long isoform IDE-Met^1^ (left panel, green) with mitochondria stained (middle panel, red), and colocalization shown (right panel, yellow). Middle row: Endogenous PYCR1 (left panel, green) with mitochondria (middle panel, red) colocalizing (right panel, yellow). Bottom row: PYCR1(left panel, green) with IDE-Met^1^ (middle panel, red) colocalizing (right panel, yellow). The insets in the upper left of each panel show expanded views of the regions indicated by white rectangles. Cell nuclei are visualized with blue DAPI stain. The red IDE secondary antibody also localized to nuclei, giving rise to the purple staining in two panels of the bottom row. There is no indication of IDE-Met^1^ localization to nuclei when using other secondary antibodies, demonstrating the observed staining of the nucleus is an artifact of the secondary antibody used.

**Table 1.** Analysis of IDE, PYCR1, and mitochondria colocalization by immunofluorescence microscopy.

| colocalization pair | Pearson's correlation coefficient<sup>1</sup> | Manders' coefficient M1<sup>2</sup> | Manders' coefficient M2<sup>3</sup> |
| --- | --- | --- | --- |
| PYCR1:mito | 0.71 | 0.76 | 0.82 |
| IDE:mito | 0.78 | 0.91 | 0.85 |
| PYCR1:IDE | 0.69 | 0.88 | 0.78 |

<sup>1</sup>Pearson's correlation coefficient = $\sum_i (I1_i - \langle I1 \rangle)(I2_i - \langle I2 \rangle) / (\sum_i (I1_i - \langle I1 \rangle)^2 \times \sum_i (I2_i - \langle I2 \rangle)^2)^{1/2}$ , where $I1_i$ and $I2_i$ are the intensities in pixel $i$ for the two color channels, and $\langle I1 \rangle$ and $\langle I2 \rangle$ are the mean intensities of the two color channels .
<sup>2</sup>Fraction of first colocalization partner fluorescence intensity located in pixels where the second partner intensity is above a threshold [54], which was determined by the Costes procedure [44].
<sup>3</sup>Fraction of the second colocalization partner fluorescence intensity located in pixels where the first partner intensity is above a threshold [54], which was determined by the Costes procedure [44].

### PYCR1 interacts with IDE *in vitro* and modulates its activity

We next set out to determine a direct functional interaction between PYCR1 and IDE. We thus looked at the effect of purified recombinant PYCR1 on purified recombinant IDE activity using the synthetic fluorogenic substrate Abz-Gly-Gly-Phe-Leu-Arg-Lys-His-Gly-Gln-EDDnp (Abz substrate). The PYCR1 cDNA we obtained from Addgene contains a 19 amino acid deletion at its C-terminus (residues 1-300, PYCR1^300^) rather that the full C-terminal extension of the wild-type enzyme. A His tagged, full length PYCR1 (PYCR1^319^) was therefore generated and purified as described in Methods. The PYCR1^319^ protein eluted from a molecular sieve column as two oligomeric forms of approximate molecular weights of ∼400 kDa and ∼200 kDa, which correspond to the molecular forms of PYCR1 previously described [45]. Each molecular form gave the expected subunit band of ∼34 kDa [45] by gel electrophoresis. Full length PYCR1^319^ increased IDE activity in a linear fashion **(Fig 6 left)**, not saturating IDE under the conditions tested.

**Fig 6.**
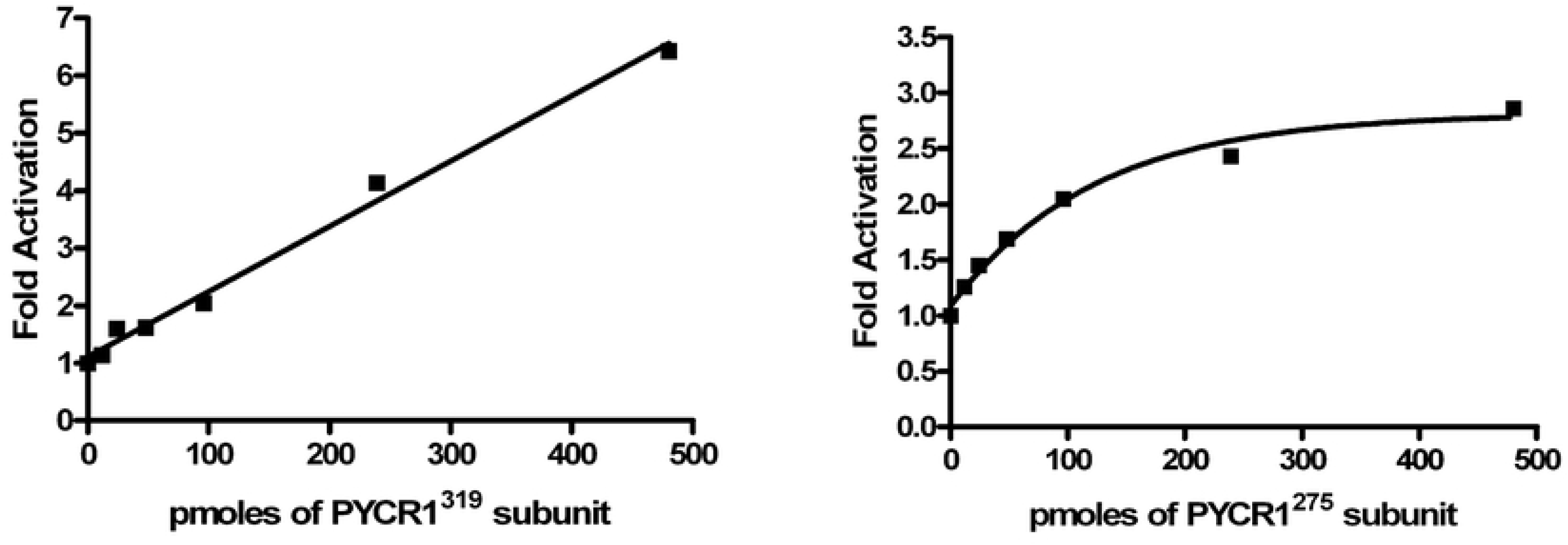
Activation of IDE by PYRC1. Shown is the increase in IDE activity produced by added PYRC1 using a small fluorogenic peptide substrate. Left: Resultant activity of IDE when increasing amounts of full-length PYRC1 (PYRC1^319^) was added to IDE. Right: Resultant activity of IDE when increasing amounts of truncated PYRC1 (PYRC1^275^) was added to IDE.

Biolayer interferometry confirms a direct interaction between IDE and PYCR1. With PYCR1^300^ bound to the biosensor, IDE at different concentrations shows well defined on and off interactions characteristic of specific binding (**Fig 7**). Global fitting of the on and off rates across three IDE concentrations gives an apparent dissociation constant of 3.0 µM (95% CI: 2.89-3.11 µM), which is well within the range of affinities for transient protein-protein interactions [56–58]. Dynamic light scattering (DLS) measurements also support direct binding of IDE and PYCR1 (see **Fig 7**). Using a globular protein model, the DLS signal from an IDE sample alone fits with a molecular weight of 276 kDa (mean hydrodynamic radius of 6.6 nm, percent polydispersity of 8.2), most consistent with the expected dimer [14]. PYCR1 fits with a molecular weight of 177 kDa (mean hydrodynamic radius of 5.4 nm, percent polydispersity of 5.2), consistent with a pentamer, which has been reported for PYCR1 [45]. (Both samples also have much higher molecular weight peaks, modeled at ∼2500 kDa, which are likely due to incomplete removal of dust particles or aggregated protein.) The mixture of IDE and PYCR1 fits with a molecular weight of 477 kDa (mean hydrodynamic radius of 8.0 nm, polydispersity index of 15.9), indicating complex formation. The modeled molecular weight is most consistent with a dimer of IDE binding a pentamer of PYCR1. It is interesting to note that the mixed sample also showed a small molecular weight species of approximately monomer PYCR1 molecular weight, perhaps suggesting some disruption of the pentamers.

**Fig 7.**
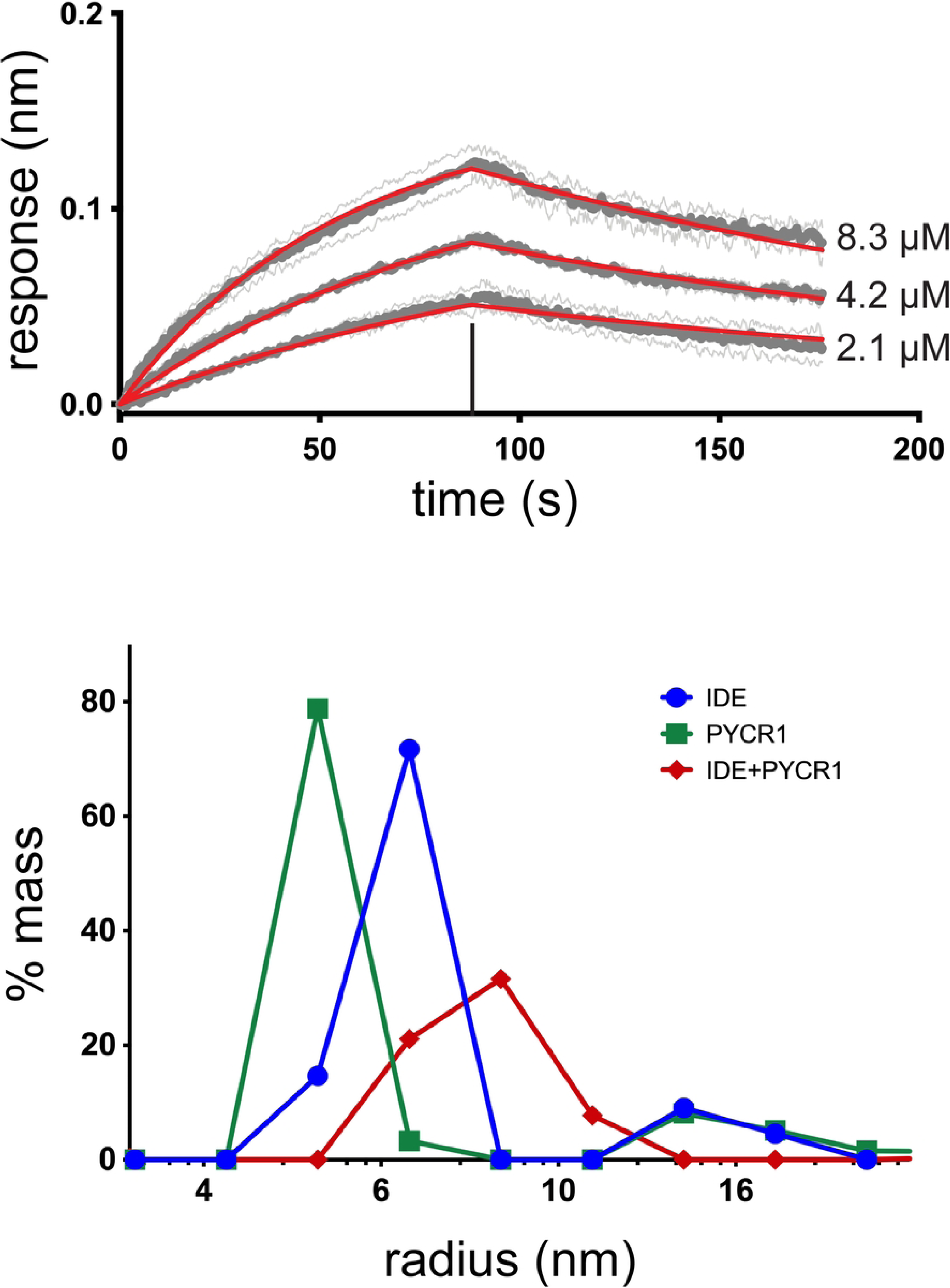
Binding interaction of IDE and PYCR1. The top panel shows biolayer interferometry association and dissociation curves for immobilized PYCR1^300^ at the three IDE concentrations indicated. The vertical black line indicates the time of conversion from association to dissociation. Triplicate averages from independent experiments are shown as dark gray circles, and fitted curves where k_on_ and k_off_ were refined globally for all three IDE concentrations are in red. Light gray lines flanking the curves and data indicate plus and minus standard deviations for each point of the triplicate data. Global refinement gave a k_on_ value of 1593 M^-1^s^-1^ (95% CI: 1541-1646 M^-1^s^-1^) and a k_off_ value of 0.0048 s^-1^ (95% CI: 0.0047-0.0049 s^-1^). Modeling of dynamic light scattering for IDE alone (blue), PYCR1 alone (green), and a mixture of the two proteins (red) is represented in the bottom panel. The semilog plot shows estimates of percent mass in the sample versus modeled hydrodynamic radius of the solute protein or proteins.

### PYCR1 makes more than one type of interaction with IDE

Using AlphaFold 2 [59], we generated a model for the interaction of PYRC1^319^ and IDE (**Fig 8)**. This model predicts the C-terminal unstructured region of PYRC1 facilitates the interaction between PYRC1 and IDE by interacting in the inner chamber of IDE in a manner similar to a bound peptide [14]. Similar C-terminal interactions were predicted for our original construct, PYCR1^300^, either as a monomer or a dimer. We thus tested the effect of removing the entire C-terminal extension (residues 276-319). This form of PYCR1 (PYCR1^275^) also produced two oligomeric forms of ∼400 and 200 KDa, both composed of the ∼34 kDa subunit as shown by SDS-PAGE. PYCR1^275^ produced a significant increase in IDE activity using the synthetic Abz substrate, however in this case a hyperbolic activation of IDE activity was observed (see **Fig 6 right**), suggesting that both the core PYCR1 and the C-terminal unstructured region interact with IDE to differently modulate the activity of the enzyme.

**Fig 8.**
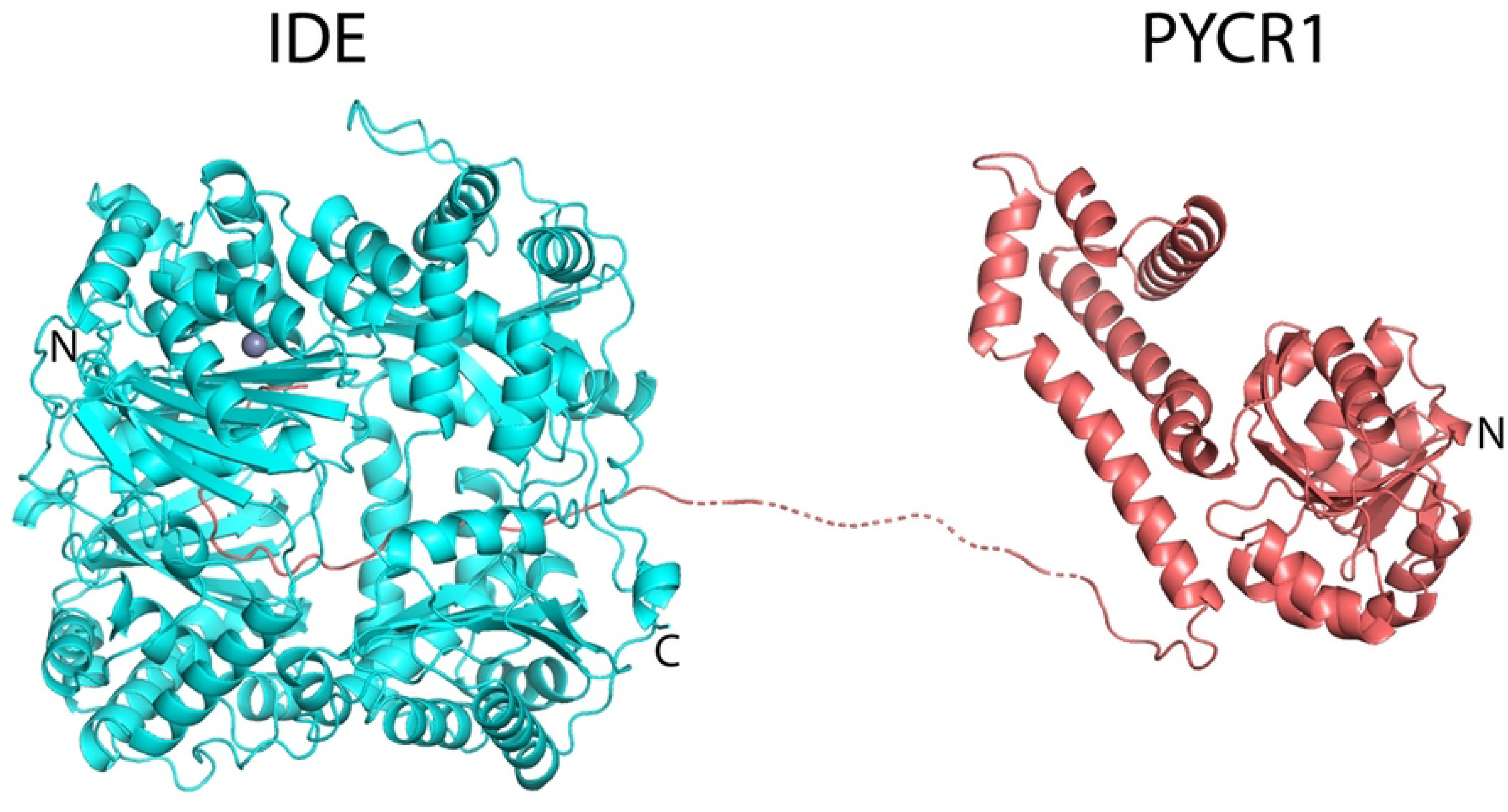
AlphaFold 2 predicted model for the interaction of PYRC1 with IDE. The C-terminal extension of PYRC1 is predicted to be inserted into the inner chamber of IDE in such a fashion that it can lead to IDE activation.

With the large peptide insulin, a known physiological substrate of IDE, PYCR1^319^ did not exhibit an effect on IDE activity nor did PYCR1 with the entire C-terminal extension removed (PYRC1^275^), but with bradykinin, a smaller (9 amino acid) physiological substrate, a maximal 4-fold increase in peptide cleavage by PYCR1^319^ was observed (**Table 2**). Both the 400 kDa and the 200 kDa oligomeric forms of PYRC1^319^ produced the same effect.

**Table 2.** PYCR1^319^-dependent increase in the cleavage of bradykinin by IDE.^1^.

|  | Fold Increase by the 400 kDa form of PYRC1 | Fold Increase by the 200 kDa form of PYRC1 |
| --- | --- | --- |
| IDE + bradykinin | 1.0 | 1.0 |
| IDE + bradykinin + PYRC1 (5.0 molar equivalents) | 1.8 | 1.9 |
| IDE + bradykinin + PYRC1 (10 molar equivalents) | 3.1 | 2.7 |

<sup>1</sup>IDE (0.25 ug) was incubated with 25 μM bradykinin for 10 min at 37°C in 50 mM Tris buffer pH 7.4 and when indicated PYCR1. The rate of hydrolysis of bradykinin was determined by measuring the decrease in the bradykinin peak area as determined by HPLC

The AlphaFold model indicates that PYCR1^319^ was bound in such a fashion that it could serve as an activator of IDE by producing a conformational change like the binding of peptide activators to IDE [14, 60]. Such was not the case for PYCR1^275^. AlphaFold 2 did not predict a significant interaction between IDE and PYCR1^275^, and the mechanism of activation by this shorter form of PYCR1 remains to be determined.

We considered whether the IDE-PYRC1 complex could produce a reciprocal activation of PYRC1 activity. Using the activity assay described by Meng et al. [46], in which PYRC1 catalyzes the oxidation of thioproline with the concomitant reduction of NADP, we found IDE had no effect on its activity up to ratio of 20 IDE to 1 PYRC1.

## Discussion

We report an intracellular protein activator of IDE, namely pyrroline-5-carboxylate reductase 1 (PYCR1). PYCR1 is most noted for converting pyrroline-5-carboxylate to proline in an NAD(P)H dependent reaction, however it has also been reported to interact with the small B subunit of ribonucleotide reductase and modulate cellular oxidative stress [61].

Evidence that PYCR1 is an IDE binding protein includes the identification of PYCR1 in pull-down experiments from three different cell lines, the demonstration of a direct interaction of IDE and PYRC1, and the finding that PYCR1 affects the activity of IDE. That a partially truncated form, PYCR1^300^, produces the same effect as full-length PYCR1^319^, while PYCR1^275^, with its C-terminal extension completely deleted, acts differently, indicates that there are two types of interactions between these proteins: one mediated by the C-terminal unstructured region of PYCR1 and one mediated by the well-folded, globular portion of the enzyme. The models we generated using AlphaFold would suggest that there is an insertion of the unstructured C-terminal residues (particularly residues 276-300) into the inner chamber of IDE. Binding in this manner may activate the enzyme by weakening the interface between the two halves of IDE, shifting the conformational equilibrium toward the open form of the enzyme [14, 60]. However, it is not just this interaction that is responsible for the binding of IDE and PYCR1, since a form of PYCR1 in which these C-terminal residues were deleted also interacts with IDE, producing a smaller, but significant, effect on activity. Unfortunately, we were unable to model this interaction using AlphaFold.

It is unlikely that PYCR1 regulates the cytosolic pool of IDE since it is localized to mitochondria. However, Selkoe and coworkers [23] described a mitochondrial pool of an IDE isoform that includes residues 1-41, and this finding has been supported by others [24–26]. We show here that the long form of IDE colocalizes with PYCR1 in mitochondria and therefore postulate that this pool of IDE is regulated by PYCR1. The function of mitochondrial IDE has not been well established. However, it has been shown that IDE can remove N-terminal mitochondrial targeting sequences from proteins [23, 62, 63], and IDE may play a role in metabolizing mitochondrial Aβ [25]. Determining if either activity is modulated by PYCR1 in mitochondria will be an important next step in continuing efforts to fully describe IDE’s functional roles.

## Acknowledgements

The authors acknowledge the use of facilities in the University of Kentucky Center for Structural Biology. We thank Dr. K. Martin Chow of the University of Kentucky Nanobody and Protein Production Facility for help with aspects of this work.

## Supporting information

**Table S1. Proteins from an HEK cell extract co-immunoprecipitated with IDE.**

**Fig S1. Nanobody binding to IDE.** Kinetics of the D1 nanobody binding to intact IDE or the N-terminal half of IDE as indicated. Bio-layer interferometry association and dissociation data following instrument response over time are shown in blue. Non-linear fits to the data curves using a single site, one-to-one model are in black. Binding of a control nanobody not raised against IDE is also shown. Vertical red lines indicate the time of conversion from association to dissociation. Affinity values are given in the text.

**Fig S2. Comparison of co-immunoprecipitation for the three cell lines used.** Lanes showing immunoprecipitation with the Fc-anti-IDE-Nb-D1 nanobody from the Coomassie-stained SDS-PAGE gels in Figs. 1 and 2 aligned for comparison. Cell lines are indicated, and the band labels refer to the proteins listed in Table S1.

**Fig S3. Co-immunoprecipitation of pyrroline-5-carboxylate reductase (PYRC1) and glucose 6 phosphate dehydrogenase (G6PD) from HeLa IDE knockout cells.** The left pane shows a Coomassie-stained SDS-PAGE gel with cell lysates subjected to immunoprecipitation by (1) anti-IDE nanobody Fc-anti-IDE-Nb-D1, (2) control nanobody Fc-anti-PP-Nb-C9, and (3) the absence of nanobody. The right pane shows a Western blot with primary antibodies indicated. Cell lysate is shown as well as IPs by (1) nanobody Fc-anti-IDE-Nb-D1 and (2) control nanobody Fc-anti-PP-Nb-C9. Arrowheads on the blots indicate G6PD (top), and PYCR-1 (bottom). Molecular weight markers are indicated.

**Fig S4. Western blot showing IDE N-terminal half expression in HeLa IDE knockout cells.** Lysate from HeLa IDE knockout cells was run on an SDS-PAGE gel and subjected to Western blotting with an anti-IDE primary antibody. Lysate samples were loaded at total protein amounts of 20 µg (left HeLa IDE KO lane) and 40 µg (right HeLa IDE KO lane). Molecular weight markers are indicated, and purified recombinant IDE (1µg) is shown for comparison.

